# Interspecies interactions in bacterial colonies are determined by physiological traits and the environment

**DOI:** 10.1101/623017

**Authors:** Sean C. Booth, Scott A. Rice

## Abstract

Interspecies interactions in bacterial biofilms have important impacts on the composition and function of communities in natural and engineered systems. To investigate these interactions, synthetic communities provide experimentally tractable systems. Agar-surface colonies are similar to biofilms and have been used for investigating the eco-evolutionary and biophysical forces that determine community composition and spatial distribution of bacteria. Prior work has focused on intraspecies interactions, using differently fluorescent tagged but identical or genetically modified strains of the same species. Here, we investigated how physiological differences determine the community composition and spatial distribution in synthetic communities of *Pseudomonas aeruginosa*, *Pseudomonas protegens* and *Klebsiella pneumoniae*. Using quantitative microscopic imaging, we found that interspecies interactions in multispecies colonies are influenced by type IV pilus mediated motility, extracellular matrix secretion, environmental parameters and the specific species involved. These results indicate that the patterns observable in mixed species colonies can be used to understand the mechanisms that drive interspecies interactions, which are dependent on the interplay between specific species’ physiology and environmental conditions.

## Introduction

The growth and survival of bacteria in natural and engineered systems is strongly influenced by how they interact with one another [1, 2]. Interactions between community members vary considerably; some interactions facilitate growth and survival, while others inhibit growth or even result in the death of one species [3]. Both faciliatory and inhibitory interactions can be mediated by secreted products, e.g. metabolite exchange [4] and antibiotic production [5] or by contact-dependent mechanisms, e.g. adhesion [6, 7] and Type VI secretion mediated killing [8]. Regardless of the specific mechanism of interaction, the composition and function of a community is influenced by the combination and sum of the various interactions between its constitutive members [9, 10].

Studies of the interactions between community members have shown how interspecies interactions influence community-intrinsic properties [11]. While competitive interactions between individual species are expected and common [12], often due to overlap of metabolic preferences [13], cooperative interactions can also be found in larger communities which enable increased biomass production [14] or resistance to antimicrobial agents [15]. Biofilms are ideal for examining the effect of interactions on a community, as cells are in close proximity to each other and are embedded within self-secreted biofilm matrix [16, 17]. For example, biofilms have been used to show how the genetic division of labor improves pellicle formation [18] and that spatial structure influences the evolution [19], formation [20] and outcome [21] of interspecies interactions. Using a synthetic, three-species community, we have shown that co-culture biofilms exhibit increased growth by particular members and enhanced stress tolerance for the entire community [22]. This occurred despite the community members competing for the same resources and was not observed for planktonic mixed cultures, suggesting that spatial organization within biofilms is important for species interactions.

Colonies grown on a nutrient surface are an efficient system for manipulating and visualizing the spatial distribution of different microorganisms, but have mostly been used for studying eco-evolutionary dynamics [23]. They have been used to characterize various spatiotemporal aspects of microbial interactions including the co-localization of mutually auxotrophic strains of *Saccharomyces cerevisiae* [24, 25], the vertical separation of differently sized cells of *Escherichia coli* [26], the sequential range expansion of nitrate reducing *Pseudomonas stutzeri* [27], the spatial separation patterns of strains of *Vibrio cholerae* [28] or *Bacillus subtilis* [29] that differ in the quantity of extracellular matrix produced, and the proportion of different antibiotic resistant/sensitive strains of *P. aeruginosa* [30]. All of these studies have focused on different strains or mutants of a single species (intraspecies interactions), differing only in the fluorescent marker they express and targeted deletion of specific genes. However, it is also clear that this approach has considerable potential for investigating interactions between different species.

In this study, we investigated the interactions between members of a model community consisting of *P. aeruginosa* PAO1, *P. protegens* Pf-5 (formerly *P. fluorescens* [31, 32]) and *K. pneumoniae* KP-1. These species can be commonly found in soils and habitats as varied as metalworking fluid or the gut of silk moths [33, 34]. When co-cultured in flow-cell biofilms, this community exhibits properties not observed in biofilms of the individual species or liquid cultures including a relative increase in *K. pneumoniae*, sharing of resistance to sodium dodecyl sulfate allowing the sensitive species, *P. protegens*, to survive, [22] and reduced production of morphotypic variants for all species [35]. As colonies offer some advantages compared to flow-cell biofilms, including higher throughput and the ability to image the entire colony, here we assessed interactions between each possible pair of these species by co-culturing them in a colony biofilm model. We determined whether these interactions were beneficial or detrimental by quantifying the area colonized by each species and qualifying where each species was found. By manipulating the community through using mutant strains and/or altered environmental conditions, we also show how organism-specific physiology determines interaction outcomes. This approach thus has the potential to identify gene systems and environmental conditions that underlie the interactions between bacteria. This work demonstrates that co-culturing in colony biofilms is a useful tool for determining the outcome of interactions between bacteria and visualizing the emergent properties of multi-species biofilms.

## Materials and Methods

### Strains, media and growth conditions

Colonies were grown on minimal medium (48 mM Na_2_HPO_4_; 22 mM KH_2_PO_4_; 9 mM NaCl; 19 mM NH_4_Cl; 2 mM MgSO_4_; 0.1 mM CaCl_2_; 0.04 mM FeSO_4_; 2 mM glucose and 0.4% casamino acids) with either 1.5% or 0.6% agar in 24 well plates or 8 well Ibidi™ slides. Previously described strains (Table 1) [22] of *P. aeruginosa* PAO1, *P. protegens* Pf-5 and *K. pneumoniae* KP-1 constitutively expressing fluorescent protein genes inserted into the chromosome using a Tn7 transposon were recovered from freezer stocks directly onto LB agar plates and grown for 24-48 h at room temperature. Bacteria were inoculated into liquid medium from agar plates (because they are highly mucoid, wild-type *K. pneumoniae* strains were homogenized by passing repeatedly through a 30 G needle), the OD_600_ was standardized to 0.1 and 1/100 dilutions were used directly or mixed 1:1 and 0.5 µL was spotted at the center of each well. Spots were dried for 30 min in a laminar flow cabinet, plates were sealed with parafilm and incubated at room temperature. To confirm that mixtures were equal, colony forming units (CFUs) were determined for the inoculum for each species/strain. If CFU counts indicated that the ratio of two mixed species/strains was more than 10 to 1 the sample was not used. For co-culture colonies, only combinations of CFP or YFP labelled strains with dsRed or mCherry were used, with the exception of KP-1-YFP and Pf-5-CFP, due to issues of overlapping excitation and emission spectra. When multiple combinations were possible (e.g. KP-1-dsRed with KP-1-CFP or KP-1-YFP) both combinations were treated as equivalent i.e. the combination of dsRed with CFP was considered a replicate of combinations of dsRed with YFP.

**Table 1:** Species and strains used in this study.

| Species and strain | Genotypic and phenotypic characteristics | Tn7 chromosomal fluorescent protein | Source or reference |
| --- | --- | --- | --- |
| *P. aeruginosa* PAO1 | Wild-type | YFP, mCherry | ATCC BAA-47 [19], this study |
| *P. aeruginosa* PAO1 Small colony variant (SCV) | *pilT* Thr210 frameshift, twitching defect | YFP | [32] |
| *P. protegens* Pf-5 | Wild-type | CFP, mCherry | ATCC BAA-477 [19], this study |
| *K. pneumoniae* KP-1 | Wild-type | dsRed, CFP, YFP | [19] |
| *K. pneumoniae* KP-1 Non-mucoid variant (NMV) | UDP-glucose lipid carrier transferase Phe123Ile, matrix secretion defect | dsRed | [32] |

### Image acquisition

Colonies were imaged on a Zeiss Axio Observer.Z1 inverted (24 well plates) or Imager.M2 upright (8 well slides) microscope using an HXP 120 C halogen light source. On the Z1 a 5X/0.16 objective was used and a 10X/0.3 objective on the M2. The filter sets (excitation/emission) used were: Zeiss Set 10 for CFP (450-490/515-565), Set 46 for YFP (490-510/520-550) and Set 31 (550-580/590-660) for dsRed and mCherry on the Z1. Set 43 (520-570/565-630) was used for dsRed and mCherry on the M1. Exposure times and focus were manually optimized for each sample then colonies were imaged using ‘Tilescan’ mode in Zeiss Zen (Blue edition) with 10% overlap. Twelve-bit images were captured using an AxioCamMRm3. Shading correction was applied using precaptured profiles of fluorescent slides. Images from both microscopes were treated equivalently in downstream processing.

### Image Presentation

Representative images from each experimental combination were selected for presentation. Images were processed for viewing to improve contrast and remove artefacts, but only raw images were used for quantitative analysis. Images were corrected for flat-field illumination using BaSiC [36], stitched together in ImageJ [37, 38] and then the brightness was scaled manually for each channel in each image. In cases when there was high background signal, the background was either subtracted manually by segmenting the colony area or by using the background subtraction tool.

### Image Analysis

For full details, see the supplementary material. Briefly, tilescan images stitched together using Zeiss Zen were imported into R and processed using the package ‘imager’ [39]. Each channel was normalized to a range of 0 to 1 and any background signal was subtracted. To segment the colony from the background, a mask of the colony was generated by taking the sum of all channels (including the inverted brightfield) then thresholding this image using either “IJDefault” or “triangle” in the R package ‘auto_thresholdr’ [40]. To determine the amount of overlap between the two strains within a colony, the ratio between channels was calculated at every pixel by subtracting one channel from the other and dividing them by the sum of both ((c1-c2)/(c1+c2)). Thus, values can range from 1/-1 indicating only the presence of one strain, to 0, indicating an exactly equal mix of the two strains. The proportion of mixed pixels was then calculated by dividing the number of bins in the range −0.25:0.25 by the total number of pixels within the colony as defined by the segmentation. We then attempted to determine the area occupied by each strain within the colony by thresholding each fluorescent channel with several different methods (including Li, Huang and triangle), then manually selecting the most accurate method. Due to the manual intervention required to pick an appropriate threshold, which varied substantially between images, the area of each strain was instead manually determined in ImageJ. For all conditions a minimum of 6 biological replicates were analyzed.

### Statistics

Analysis of variance, Tukey’s honest significant difference post-hoc test and multiple-testing p-value adjustment were used to determine if there were significant differences in colony area between single and dual species colonies.

## Results

### Monospecies Colonies

To compare patterns of interactions with previously published studies, we first examined intraspecies interactions by growing colonies consisting of two strains of the same species (Figure 1). Qualitative inspection of the colonies after 24 and 48 h of growth indicated that each species differed in how they separate into sectors. *P. protegens* (Figure 1B,E) and *K. pneumoniae* (Figure 1C,F) showed clear segregation between strains, although the shape of the borders differed. For *P. aeruginosa* (Figure 1A,D), separation into sectors was only apparent at 48 h and the borders between strains were less distinct than for the other species.

**Figure 1:**
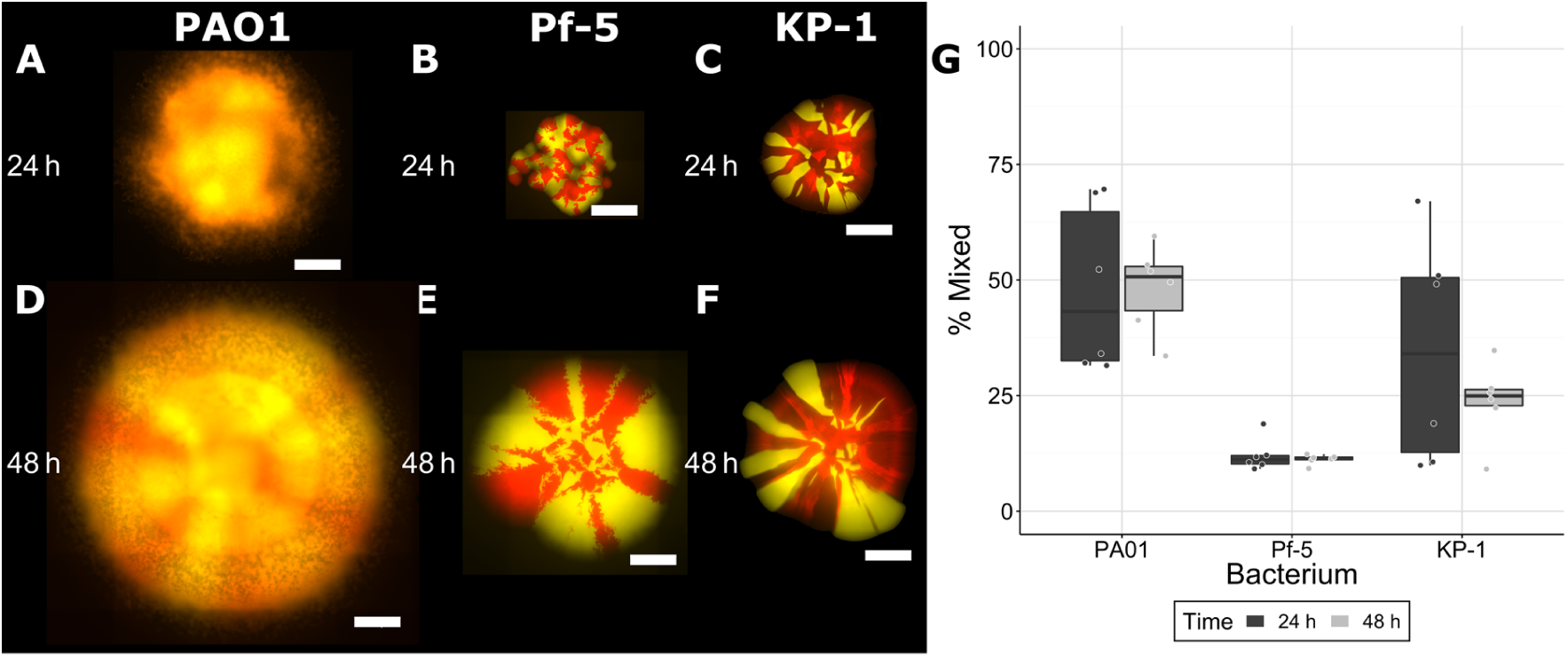
Colonies consisting of *P. aeruginosa* PAO1 (A, D), *P. protegens* Pf-5 (B, E) or *K. pneumoniae* KP-1 (C, F). Each species was labeled separately with two different fluorescent protein genes and are presented here as red or yellow (false colouring). Strains were mixed 1:1 (OD_600_) at the time of inoculation and colonies were imaged after 24 and 48 h of growth. Scale bars indicate 1000 µm. Overlap between strains was calculated from raw images as the percentage of mixed pixels (G). Boxplots are presented in Tukey style.

To better characterize the differences in colony growth, quantitative analyses were performed by determining the percentage of pixels with similar relative signal within each colony (Figure 1, G). Around 50% of pixels were mixed in the *P. aeruginosa* colonies compared to 25% or less in the other two species after 48 h of growth. Even at the leading edge of *P. aeruginosa* regions that appeared to consist of a single strain, cells of both colours were present and were well mixed (Supplementary Video 1). Movement and mixing of the two strains was visible 500 µm inward from the colony edge (Supplementary Video 2). For all three species, there was no change in mixing between 24 and 48 h.

### Dual Species Colonies

In contrast to monospecies colonies, dual species colonies differed in the area colonized and mixing of species (Figure 2). None of the combinations of species separated into clear radial sectors, as observed for the dual-strain colonies described above, and instead formed distinct patterns. *P. aeruginosa* localized around the perimeter of *P. protegens* colonies (Figure 2A,D), especially after 48 h and these species did not mix at either timepoint. While *P. aeruginosa* appeared to be restricted to the interior of the colony when cultured with *K. pneumoniae* (Figure 2B,E), closer inspection of brightfield images (Supplementary Figure 1) showed that *P. aeruginosa* was also present around the outside perimeter of *K. pneumoniae*, as visible with its distinct lattice morphology. This combination changed from unmixed (~25%, though variable) at 24 h to mixed at 48 h (consistently 50%). When grown with *K. pneumoniae*, *P. protegens* grew around the outer edge of the colony (Figure 2C,F) and these species were also partially mixed (~40%) at 48 h. The amount of area covered by each strain in the dual-species colonies was compared to the area covered by single species colonies (Figure 2, H, Supplementary Table 1). At 24 h, the area covered by *P. aeruginosa* (~8 mm^2^) was lower when co-cultured with *P. protegens* (~2 mm^2^) and was similarly reduced at 48 h, from ~70 mm^2^ (monospecies) compared to ~35 mm^2^ (dual species). In contrast, there was no difference in the area covered by *P. protegens* when co-cultured with *P. aeruginosa*, although it was significantly decreased (from ~4 to ~1.5 mm^2^) at 24 h, but not 48 h, with *K. pneumoniae*. The area covered by *K. pneumoniae* increased when co-cultured with *P. aeruginosa* from ~5 to ~10 mm^2^ at 24 h and ~12 to ~40 mm^2^ at 48 h, but was not affected by *P. protegens*.

**Figure 2:**
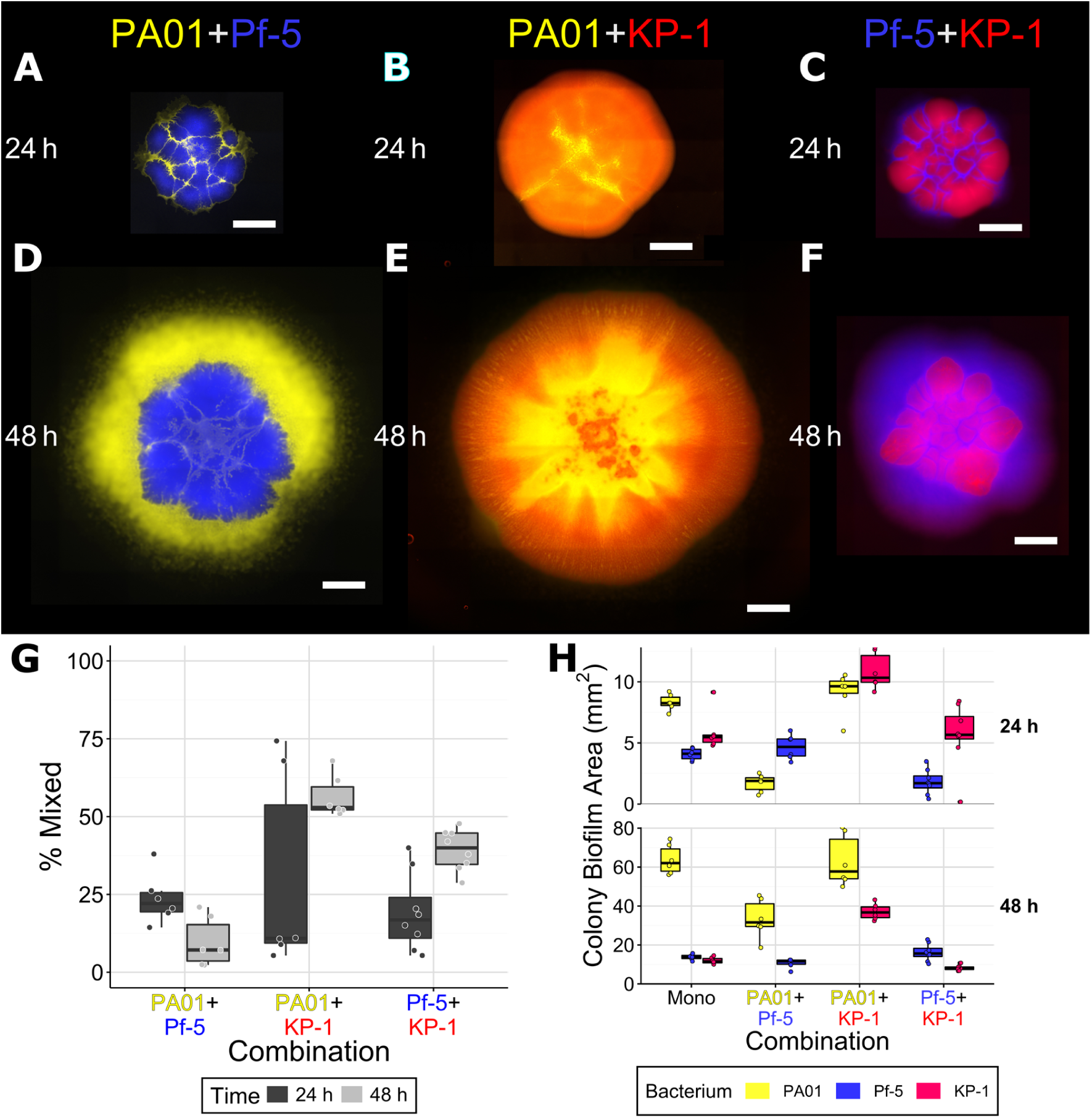
Dual species colonies of *P. aeruginosa* PAO1 and *P. protegens* Pf-5 (A, D), *P. aeruginosa* PAO1 and *K. pneumoniae* KP-1 (B, E), and *P. protegens* Pf-5 and *K. pneumoniae* KP-1 (C, F), overlap between differently labeled strains (G) and the area covered by individual species in colonies (H). Each species was labeled with different fluorescent protein genes and are presented here as yellow (*P. aeruginosa* PAO1), blue (*P. protegens*) and red (*K. pneumoniae*). Strains were mixed 1:1 (OD_600_) at the time of inoculation and colonies were imaged after 24 and 48 h of growth. Scale bar indicates 1000 µm. Boxplots are presented in Tukey style.

### Co-cultures with a *P. aeruginosa* Small Colony Variant

To investigate the effect of the type IV pilus (TFP) of *P. aeruginosa* in colonies, a small colony variant (SCV) was cultured in dual species colonies with the parental strain, *P. protegens* and *K. pneumoniae*. Based on whole genome sequencing, we previously determined that this SCV differs from the wild-type only by a single mutation in *pilT* [35], preventing pilus retraction and therefore TFP motility [41, 42]. Colonies formed with the SCV differed from those with wild-type *P. aeruginosa* and the mixing of strains/species also differed (Figure 3). Both *P. aeruginosa* (Figure 3A,D) and *P. protegens* (Figure 3B,E) formed a perimeter around the SCV, but there was very little (≤10%) mixing of strains/species. The SCV only mixed slightly more with *K. pneumoniae* (Figure 3C,F). The SCV colonized less area than wild-type *P. aeruginosa, ~*12 mm^2^ compared to ~70 mm^2^, respectively, after 48 h (Figure 3, H). In dual species colonies, the area covered by the SCV was reduced to ~2 mm^2^ at 48 h in the presence of *P. protegens*. When grown with *K. pneumoniae*, the SCV covered less area at 24 h but this was not apparent at 48 h. The SCV did not affect the area covered by the other strains and in contrast to when the parental strain of *P. aeruginosa* was present, the SCV did not increase the area covered by *K. pneumoniae*.

**Figure 3:**
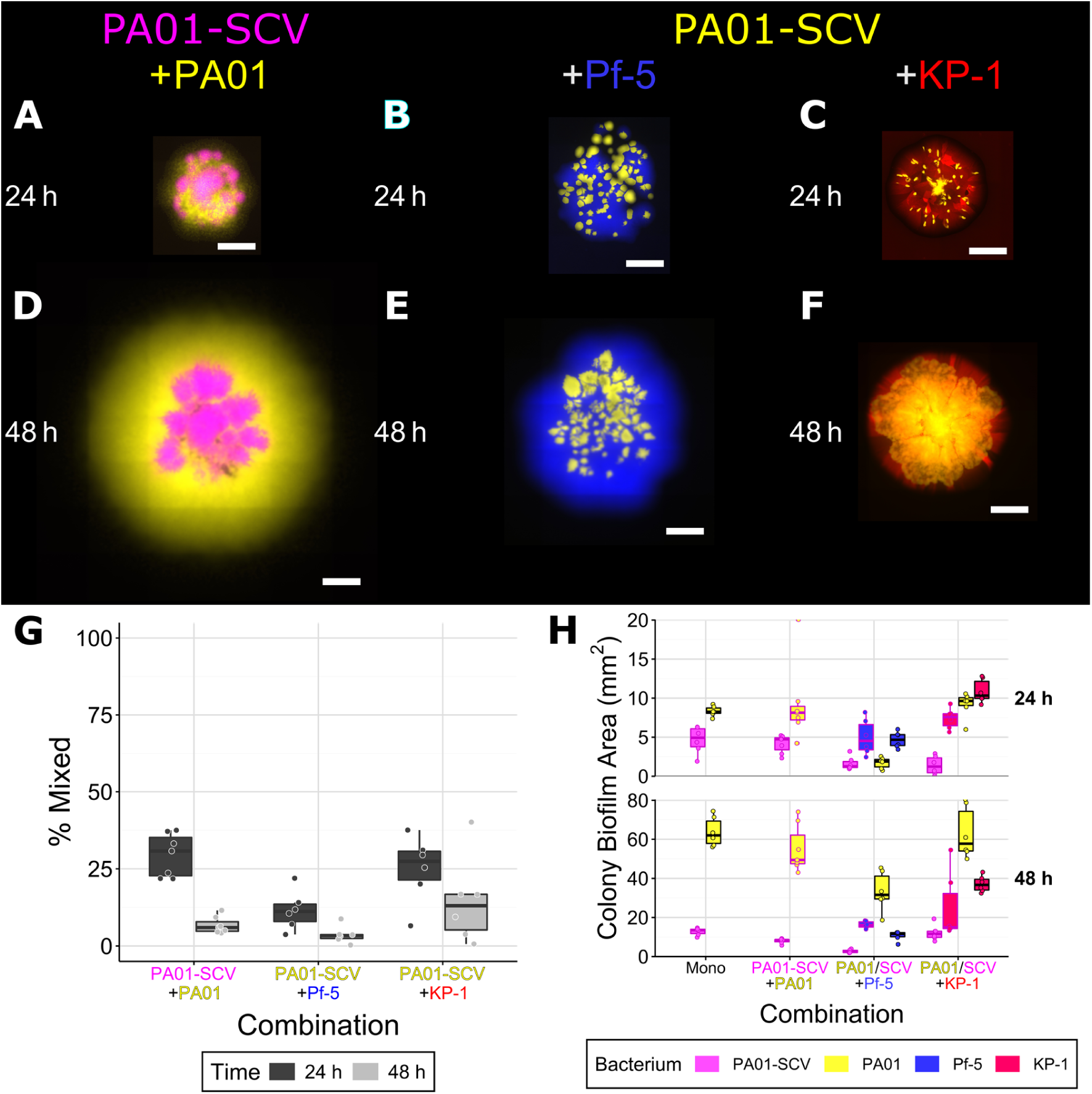
Co-culture colonies consisting of *P. aeruginosa* PAO1 and *P. aeruginosa* PAO1 small colony variant (SCV) (A, D), *P. aeruginosa* PAO1 SCV and *P. protegens* Pf-5 (B, E) and *P. aeruginosa* PAO1 SCV and *K. pneumoniae* KP-1 (C, F), overlap between differently labeled strains (G), and the area covered by individual species/strains (H). Each species was labeled with different fluorescent protein genes and are presented here as yellow (*P. aeruginosa* wild-type when paired with the SCV, SCV when paired with the other two species), magenta (*P. aeruginosa* SCV), blue (*P. protegens*) and red (*K. pneumoniae*). Strains were mixed 1:1 (OD_600_) at the time of inoculation and colonies were imaged after 24 and 48 h of growth. Scale bar indicates 1000 µm. Boxplots are presented in Tukey style. For the area boxplots (H) combinations of *P. aeruginosa* with *P. protegens* or *K. pneumoniae*, boxes outlined in black represent the wild-type (same data as Figure 2H), outlines in magenta represent combinations with the SCV.

### Co-cultures with a Non-Mucoid *K. pneumoniae* Variant

To investigate the effects of extracellular matrix (ECM) production by *K. pneumoniae*, a non-mucoid variant (NMV) was cultured in dual strain/species colonies with the parental strain, *P. aeruginosa* and *P. protegens*. This previously characterized NMV has a mutation within a probable colanic acid synthesis gene cluster, preventing it from producing its normal, thick ECM [35]. Single-strain colonies formed by the NMV differed from wild-type *K. pneumoniae* (Supplementary Figure 2). Boundaries between sectors of the NMV were jagged instead of straight as was observed for the wild-type. Dual-species colonies with the NMV also differed as it did not mix with wild-type *K. pneumoniae* or *P. protegens* (Figure 4). When grown with *P. aeruginosa* (Figure 4B,E), mixing was low at 24 h (10-20%) and was not readily observed at 48 h. When grown as single-strain colonies, wild-type *K. pneumoniae* and the NMV each covered a similar amount of area. When the two were co-cultured together (Figure 4A,D), the NMV covered less area, ~2.5 mm^2^, compared to when grown alone, ~7 mm^2^, at 24 h, and was also reduced at 48 h (from ~12 to ~5 mm^2^). The area covered by the NMV was also decreased when paired with *P. protegens* (Figure 4C,F), however, neither of these differences persisted at 48 h. In contrast, the area covered by *P. aeruginosa* was decreased when grown in the presence of the *K. pneumoniae* NMV at 24 h but not 48 h. When paired with *P. aeruginosa*, the area covered by the NMV was not significantly increased as was observed for the parental *K. pneumoniae* strain when paired with *P. aerguinosa*.

**Figure 4:**
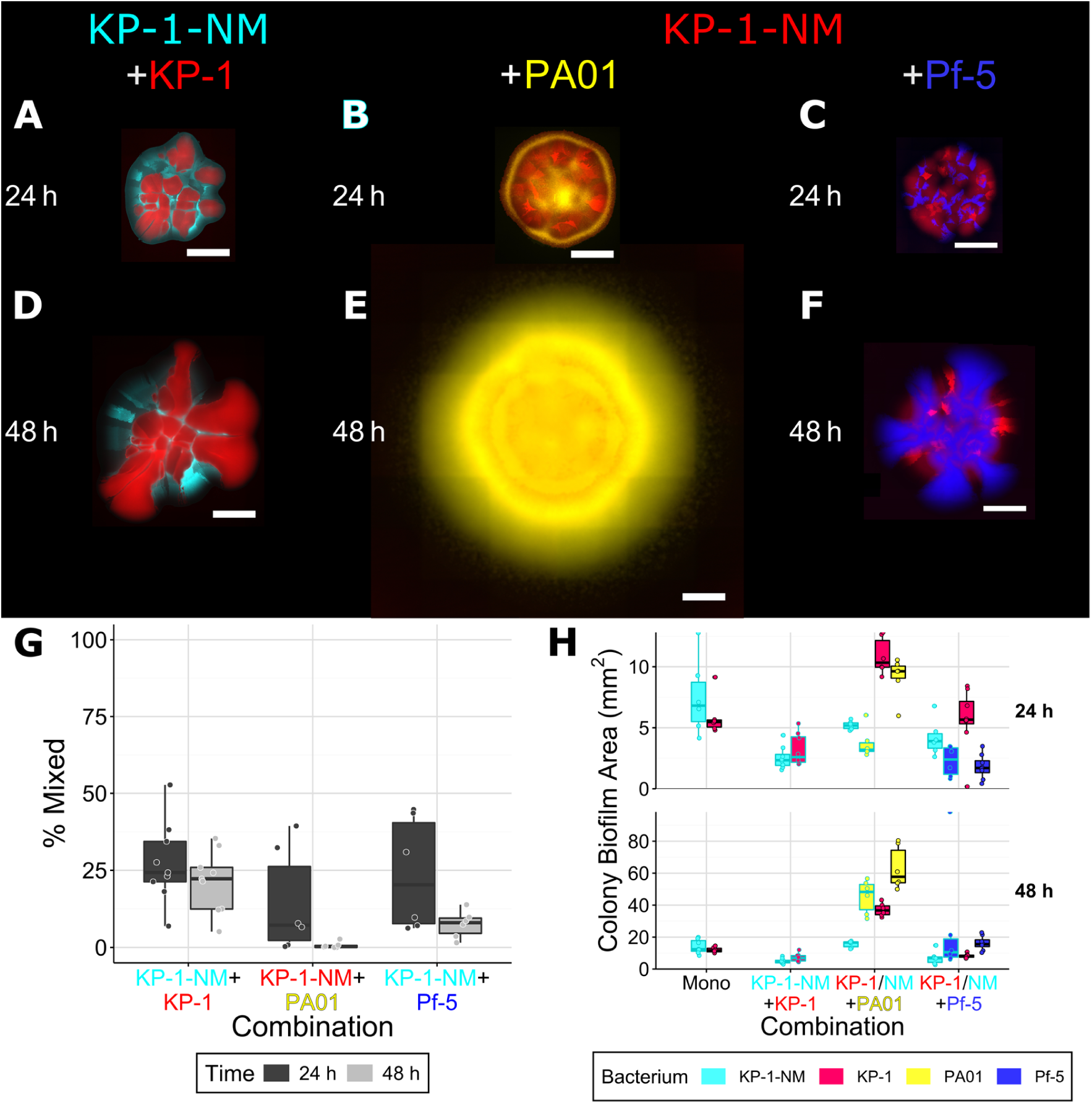
Co-culture colonies consisting of *K. pneumoniae* KP-1 with the *K. pneumoniae* KP-1 non-mucoid variant (NMV) (A, D), *K. pneumoniae* KP-1 NMV with *P. aeruginosa* (B, E), and *K. pneumoniae* KP-1 NMV with PAO1 *P. protegens* Pf-5 (C, F), overlap between differently labeled strains (G) and the area covered by individual species/strains (H). Each species was labeled with different fluorescent protein genes and are presented here as yellow (*P. aeruginosa*), blue (*P. protegens*), red (*K. pneumoniae* wild-type when paired with the NMV, *K. pneumoniae* NMV when paired with the other species) and cyan (*K. pneumoniae* NMV). Strains were mixed 1:1 (OD_600_) at the time of inoculation and colonies were imaged after 24 and 48 h of growth. Scale bar indicates 1000 µm. Boxplots are presented in Tukey style. For the area boxplots (H), combinations of *K. pneumoniae* with *P. aeruginosa* or *P. protegens*, boxes outlined in black represent the wild-type (same data as Figure 2, H), outlines in cyan represent the NMV.

### Colonies Cultured on 0.6% Agar

The percentage of agar influences motility, as lower percentages enable swarming and/or swimming motility and affect the colony growth of highly mucoid strains [28]. Culturing on a lower percentage of agar was thus expected to affect interactions between the wild-type strains of *P. aeruginosa* and *K. pneumoniae* and their non-motile (SCV) and non-mucoid (NMV) variants. Colonies consisting of wild-type and mutant strains grown on 0.6% agar had altered morphologies (Figure 5) and areas colonized (Table 2), but those consisting of two differently labeled wild-type strains did not. Additionally, the wild-type strains did not differ in how they separated into sectors (Supplementary Figure 3). When the *P. aeruginosa* SCV was co-cultured with its parental strain (Figure 5A,C), both strains expanded outwards together, but were less mixed (~22% at 48 h) compared to the two differently coloured wild-type *P. aeruginosa* strains (~55% at 48 h, Figure 1G), which was similar to that observed on 1.5% agar (50%, Figure 1G). When the *K. pneumoniae* NMV was co-cultured with the wild-type (Figure 5B,D), the two strains showed little mixing, however by 48 h the wild-type had expanded much more than the NMV (Figure 5, D). When two wild-type *K. pneumoniae* strains were grown together, they mixed comparably on both percentages of agar (~25%, Figure 5E, Figure 1G). Colony area was significantly higher for the *P. aeruginosa* SCV on 0.6% compared to 1.5% agar at 48 h (~20 vs ~10 mm^2^, Figure 5F). This was significantly lower than the wild-type on 0.6% agar (~50 mm^2^), which was also significantly decreased compared to on 1.5% agar (~70 mm^2^). When cultured with the SCV on 0.6% agar, wild-type *P. aeruginosa* area was significantly reduced at both timepoints (from 45 to 22 mm^2^ at 48 h). The *K. pneumoniae* wild-type colonized a significantly increased area on 0.6% agar (~27 mm^2^) compared to the NMV (~10 mm^2^) and the wild-type on 1.5% agar (~11 mm^2^). When they were co-cultured, the area colonized by wild-type, but not NMV *K. pneumoniae*, was significantly lower after 48 h.

**Table 2:**
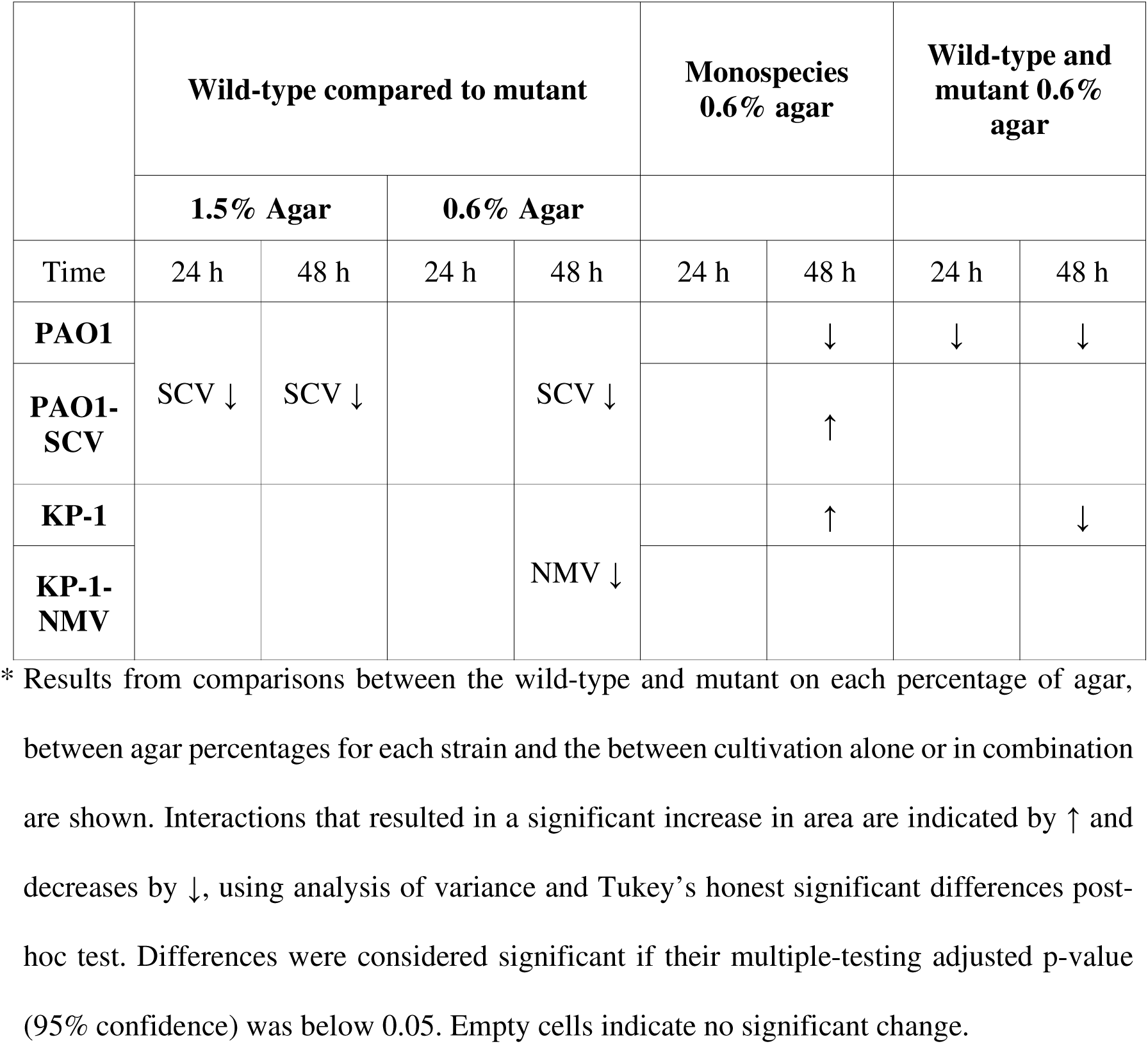
Comparison of effects of 0.6% agar on colonies of *P. aeruginosa*, *P. aeruginosa* small colony variant (SCV), *K. pneumoniae* and *K. pneumoniae* non-mucoid (NM) variant.

**Figure 5:**
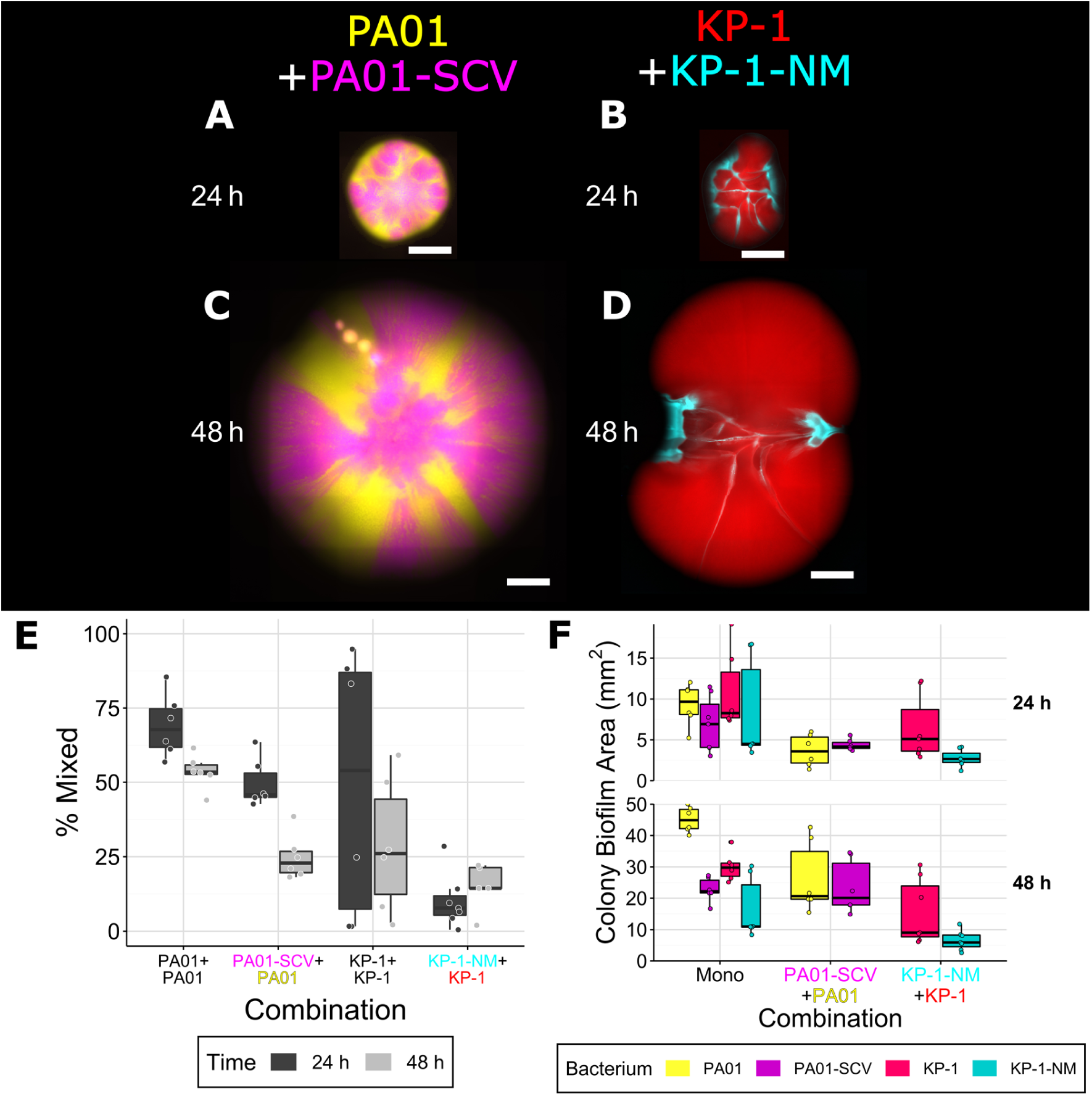
Co-culture colonies on 0.6% agar consisting of *P. aeruginosa* PAO1 with *P. aeruginosa* small colony variant (SCV) (A, C), and *K. pneumoniae* with *K. pneumoniae* KP-1 non-mucoid variant (NMV) (B, D), overlap between differently labeled strains (E) and the area covered by individual species/strains (F). Each species was labeled with different fluorescent protein genes and are presented here as yellow (*P. aeruginosa* wild-type), magenta (*P. aeruginosa* SCV), red (*K. pneumoniae* wild-type) and cyan (*K. pneumoniae* NMV). Strains were mixed 1:1 (OD_600_) at the time of inoculation and colonies were imaged after 24 and 48 h of growth. Scale bar indicates 1000 µm. Boxplots are Tukey style.

## Discussion

### Colonies highlight differences in intraspecies interactions

For each of the three species from our model community, intraspecific competition resulted in species-specific patterns of sector formation. Separation of mixed colonies into sectors has been attributed to genetic drift at the leading edge of the colony, as despite having equal fitness, stochastic effects result in new territory being colonized by either one strain or the other, but not both [43]. Here we show that the shape of the boundaries between sectors depends on the bacterium’s physiological traits. The separation of wild-type *K. pneumoniae* strains into sectors with straight boundaries resembled that of *Saccharomyces cerevisiae* [25], whereas the jagged boundaries of *P. protegens* were more similar to those of *P. stutzeri* [27], simulated rod-shaped cells [44] or *E. coli* [43]. In the case of *E. coli*, the formation of fractal boundaries between sectors has been explained by the anisotropic forces of cell division and growth causing unlinked chains of rod-shaped cells to buckle [45]. Thus, similar buckling likely causes the formation of jagged edges between boundaries of the wild-type *P. protegens* strains. The jagged boundaries visible between sectors within *K. pneumoniae* NMV colonies (Supplementary Figure 2) indicate that its secreted extracellular matrix causes the straight boundaries between sectors observed in the wild-type. Colonies of *P. aeruginosa* grown on nutrient rich LB medium were previously shown to be well mixed [46], similar to our observations here using minimal medium. The observation that *pilB* mutants segregate into clear sectors [46] and the resemblance of videos of the edge of *P. aeruginosa* colonies (Supplementary Video 1) to those showing TFP motility [47] indicates that this motility likely enabled the mixing of the *P. aeruginosa* wild-type strains.

### Interspecies interactions differ from intraspecies interactions

Previous studies have focused on colonies of either identical, but differently labeled strains of the same species, or strains with specific genetic modifications, whereas here, we investigated interactions between different species. While the patterns for our single strain data resemble previous work, it is clear that patterns of species distribution in mixed species colonies differ markedly. Thus, the data presented here show how the different physiological traits of the three studied bacteria determine community composition and spatial distribution in colonies.

We have assessed the three pairwise interactions tested here using the notation of Momeni et al. [24], where A [~,~] B indicates a neutral interaction and A[↑,↓]B indicates a relationship where A benefits and B is negatively affected (Table 3). The spatial distribution patterns of *P. aeruginosa* with *P. protegens*, and *P. protegens* with *K. pneumoniae* are similar as one strain expanded outward faster and surrounded the other. Although the area covered by the inner strains was not reduced compared to when they were grown alone, both interactions negatively affect the inner strain as it no longer had equal access to space/nutrients. For the first pair, the area covered by the outer strain, *P. aeruginosa*, was significantly decreased, indicating that both strains experienced negative outcomes from the interaction, so *P. aeruginosa* [↓,↓] *P. protegens*. For the second pair, *P. protegens* was the outer strain but its area was not reduced so, *P. protegens* [~,↓] *K. pneumoniae*. In the third case, when *P. aeruginosa* was paired with *K. pneumoniae*, *P. aeruginosa* did not differ in the amount of area covered but the area of *K. pneumoniae* was significantly increased, so, *P. aeruginosa* [~,↑] *K. pneumoniae*. Consistent with our previous work comparing planktonic and biofilm growth modes, these results differ from our observations in planktonic culture where *P. protegens* outgrew the other two species by between 10-100 fold and *P. aeruginosa* and *K. pneumoniae* equally reduced each other’s growth by ~100 fold [22].

**Table 3:** Comparison of dual species interactions for *P. aeruginosa*, *P. aeruginosa* small colony variant (SCV), *P. protegens*, *K. pneumoniae* and *K. pneumoniae* non-mucoid variant (NMV).

| **Recipient**<br>**Actor** | *P. aeruginosa* | *P. aeruginosa*<br>SCV | *P. protegens* | *K. pneumoniae* | *K. pneumoniae*<br>NMV |
| --- | --- | --- | --- | --- | --- |
| *P. aeruginosa* |  |  |  | ↑ |  |
| *P. aeruginosa*-SCV | ↓ |  |  |  | ND |
| *P. protegens* | ↓ | ↓ |  | ↓ |  |
| *K. pneumoniae* |  |  |  |  | ↓ |
| *K. pneumoniae*-NMV |  | ND |  |  |  |

\* Empty cells indicate no difference between single and dual species colonies was observed, ND indicates a combination which was not determined. Interactions that resulted in a significant increase in area are indicated by ↑ and decreases by ↓ compared to the monospecies colony by analysis of variance and Tukey's honest significant differences post-hoc test. Differences were considered significant if their multiple-testing adjusted p-value was below 0.05 for 48 h comparisons.

The interactions between *K. pneumoniae* and the two *Pseudomonas* species were starkly different. When it was grown with *P. aeruginosa* it colonized more than twice as much area than it could alone, indicating a strong benefit from being co-cultured. *P. fluorescens* Pf0-1 has been observed to rapidly evolve a division of labour where two different mutant strains are able to make a faster expanding colony than either the wild-type or each strain individually [48]. In those mixed colonies, one strain produced the force for colony expansion through cell division while the other produced an extracellular polymer which acts as a lubricant at the >expanding edge of the colony. In our work, *K. pneumoniae* may have been taking advantage of the extracellular DNA [49] or rhamnolipid surfactants [50] produced by *P. aeruginosa* to enable colony expansion. Non-motile *E. coli* was observed to take advantage of motile *Acinetobacter baylyi*, forming intricately branched floral patterns [51]. These patterns were qualitatively dissimilar to those we observed, indicating that the mechanism by which *K. pneumoniae* takes advantage of *P. aeruginosa* motility is also likely different. This strong effect of *P. aeruginosa* on *K. pneumoniae* is consistent with our observations in flow cell biofilms where despite making up only a small (1-5%) proportion of the 3-species community, *P. aeruginosa* influences the relative proportions of both other bacteria [22]. Conversely, *P. protegens* did not assist *K. pneumoniae* but was found around the outside of *K. pneumoniae* and these two strains were also more mixed at 48 h than monospecies colonies of each species. In strains of *E. coli* engineered to have equal growth rates but different cell shapes, small cells were found to reside on top of the colonies and larger cells were below, at the agar surface, where they maintained access to nutrients obtained from an agar surface [26]. *K. pneumoniae* cells are larger than *Pseudomonas* which may have allowed *P. protegens* to overgrow it and prevent *K. pneumoniae* from expanding further. In the third case, *P. aeruginosa* was also found around the outside of *P. protegens*, but these species were not mixed. This indicates that *P. protegens* could not take advantage of *P. aeruginosa*’s motility and these species mutually exclude each other in space.

### *P. aeruginosa* twitching motility is important for interactions

Twitching motility by *P. aeruginosa* has been best studied in interstitial biofilms where cells are sandwiched between an agar surface and a glass coverslip [42, 47, 49, 52]. The colonies studied here are qualitatively similar, although the expanding front does not form as intricate lattices. Without a functional TFP, the SCV could only colonize about 20% of the area of the wild-type. When paired with wild-type *P. aeruginosa* or *P. protegens*, the SCV was outcompeted for space, indicating that TFP motility is a competitive trait. It also appears to be a cooperative trait as the area colonized by *K. pneumoniae* was not increased when paired with the *P. aeruginosa* SCV, whereas it was increased three-fold when paired with TFP motile, wild-type *P. aeruginosa*. TFP motility can also be abrogated by deleting the gene for pilin subunits, pilA [52]. It will be interesting to contrast our observations of a hyper-piliated *pilT* mutant with a non-piliated *pilA* mutant. In flow-cell biofilms, a *pilA* mutant of *P. aeruginosa* was less competitive with *Agrobacterium tumefaciens* [53], but conversely, outcompeted *Staphylococcus aureus* [54]. Motility, as a competitive trait, has been suggested to allow strains better access nutrients, to cover other organisms or to disrupt their biofilms [55], which is supported by our results. Even though *P. aeruginosa* was able to cover *K. pneumoniae* with and without a functional TFP, only the motile wild-type *P. aeruginosa* had a commensal relationship with *K. pneumoniae*, indicating that motility can also be a cooperative trait.

### *K. pneumoniae* secreted matrix is important for interactions

Self-secreted extracellular matrix is a hallmark of biofilms that influences the spatial positioning and interactions between cells within a biofilm [23]. Matrix secretion allows producing cells to better access nutrients in colonies of *P. fluorescens* [56] and for simulated cells under flow conditions [57], by excluding other cells. In flow-cell biofilms, the *K. pneumoniae* NMV outcompeted its isogenic wild-type strain, but was less fit when grown with *P. aeruginosa* and *P. protegens* [35]. In colony biofilms, the *K. pneumoniae* NMV colonized the same total area as the wild-type strain when alone, but did not mix when the two were co-cultured. Furthermore, the NMV did not colonize more area when paired with *P. aeruginosa* and was outcompeted by *P. protegens*, resulting in less area covered compared to the NMV when cultured alone. This indicates that the biofilm matrix normally produced by *K. pneumoniae* KP-1 improves interspecies, but not [58] intraspecies, competition. The mutual exclusion observed between the wild-type and NMV also indicates that the NMV cannot take advantage of the wild-type, similarly to exclusion by *B. subtilis* [59] and *V. cholerae* [60]. When co-cultured, *Pantoea agglomerans* and *B. subtilis* form colonies with structural properties not observed in either single species alone, even with a *B. subtilis* mutant that does not produce EPS, indicating that it could share the exopolysaccharide being produced by *P. agglomerans* [58]. Our results indicate that the *K. pneumoniae* NMV can not similarly make use of *Pseudomonas* matrix components.

### Agar concentration changes inter-strain interaction outcomes

It is well understood that agar concentration influences bacterial motility [61]. It also affects the ability of biofilms to extract nutrients as it determines the osmotic pressure of an environment [28]. Here we showed that lowering the agar concentration from 1.5% to 0.6% increased the area colonized by the TFP motility deficient *P. aeruginosa* SCV, though it was still less than the wild-type. However, in co-culture it colonized as much area as the wild-type and was not encircled by the parental wild-type (Table 3). Conversely, the *K. pneumoniae* NMV was less effective at colonization compared to the wild-type on 0.6% but not 1.5% agar, but was still able to compete with the wild-type when co-cultured in both situations. For the *P. aeruginosa* SCV, lowering the agar concentration likely enabled flagella-based motility that could partially compensate for the defect in twitching motility. Motility is important for colonization of roots [62], the gastrointestinal tract of Zebra fish [63] and for the persistence of uropathogenic *E. coli* [64]. Here, we showed that motility influenced intraspecies competition but the outcome depended on environmental conditions. Previously, we observed that the *P. aeruginosa* SCV completely outcompeted wild-type *P. aeruginosa* in flow cell biofilms [35]. The increased attachment provided by hyper-piliation of the SCV [42, 65] likely provides this benefit in flow cells, whereas the lack of motility was a detriment in colonies. This highlights the differences in environmental pressures between the culturing methods. Hyper-piliation also leads to aggregation [66], which may explain how the *P. aeruginosa* wild-type and SCV separated into sectors with straight boundaries on 0.6% agar. This appeared to be similar to the differently tagged strains of *K. pneumoniae*, indicating that this community morphology may not exclusively be caused by matrix secretion.

For *K. pneumoniae* (which is non-motile), lowering the agar concentration allowed the wild-type to colonize more area, which was similar to how a rugose strain of *V. cholerae* that hypersecretes extracellular matrix (ECM)formed larger colonies on lower concentrations of agar [28]. In *V. cholerae*, it was demonstrated that matrix secretion generates an osmotic pressure gradient between the agar and the colony, allowing the colony to expand by physical swelling and also drawing more nutrients out of the agar. It is likely that this is a general mechanism attributable to the biofilm matrix. In this context, the wild-type *K. pneumoniae* would be similar to the rugose *V. cholerae*, producing larger amounts of ECM, while the *K. pneumoniae* NMV and wild-type *V. cholerae* are analogous in their relatively lower amount of ECM production. The community morphology of dual strain colonies of ECM secretors and non-secretors differed between the two species. In *V. cholerae*, the hyper-secretor was encircled by the non-secretor at both 1.5 and 0.6% agar, but colonized far more area at 0.6% as the non-secretor was pushed to the outside of the colony. Conversely, the *K. pneumoniae* NMV was not displaced and even prevented spreading by wild-type *K. pneumoniae* (Figure 5, D). Similar to *V. cholerae*, the *K. pneumoniae* NMV did not benefit from the matrix secreted by wild-type *K. pneumoniae* as its area was not increased in co-culture. In *B. subtilis*, it was also observed that ECM-producing cells outcompete non-secretors [29], and to a higher degree when the humidity is higher (which is similar to lower agar concentration). Additionally, osmotic pressure generated by the ECM enhances colony spreading in this species [67]. Thus, matrix production appears to be a general strategy of bacteria that increases competitiveness in colonies.

### Conclusions

Interactions between bacteria are key for determining the composition and function of communities. Here we used colonies to investigate intra- and inter-species interactions. Using the members of our three species model community, we showed that they differ in how they interact with members of their own species due to their physiological traits: TFP motility in *P. aeruginosa* allowed populations to mix whereas ECM in *K. pneumoniae* caused straight borders between population sectors. Using mutants deficient in these traits, we showed that their impact depended on the agar concentration. These physiological traits were also important when interspecies interactions were examined. The motility of *P. aeruginosa* enabled it to outcompete *P. protegens* and to facilitate increased colonization area by *K. pneumoniae*. Importantly, the spatial distribution of species, observed for dual-species colonies did not resemble any patterns previously seen for experimental or simulated monospecies and dual-strain colonies. These experiments show that interspecies interactions differ substantially from intraspecies interactions and that co-culture colonies are a powerful way to investigate how bacterial physiology determines these interactions.

## Acknowledgments

The authors would like to thank Sujatha Subramoni for helpful advice regarding the three species community, Talgat Sailov for microscopy assistance and Diane McDougald for reading the manuscript and providing helpful comments. We would also like to thank the Singapore Centre for Environmental Life Sciences Engineering (SCELSE), whose research is supported by the National Research Foundation Singapore, Ministry of Education, Nanyang Technological University and National University of Singapore, under its Research Centre of Excellence Programme.

