## Supplementary Figures and Tables for "Interspecies interactions in bacterial colonies are determined by physiological traits and the environment"

### **Supplementary Methods**

Detailed image analysis methods: Preprocessing. Images were automatically stitched together based on the collection of the tile images, then then scaled to 1/10$^{th}$ the original size in ImageJ (v1.52, [1]) to facilitate computational analyses. Images were then were imported into R for further processing using the R package ‘imager’ [2]. Channel Normalization. Each channel was normalized for each image to a range of 0 to 1 by subtracting the minimum value from all pixels then dividing them by the original range of values (i.e. the maximum minus the minimum). Background Subtraction. Backgrounds (taken to be the mean pixel value of the corners defined as 1/20 of the image) were subtracted from images with sufficient dynamic range (mean background value smaller than total range of values), and values below zero were set to zero. Masking. The location of the colony biofilm was found by taking the sum of all channels (including the inverted brightfield) then thresholding this image using the auto_thresh function from the R package ‘auto_thresholdr’ [3] using either the “IJDefault of “Triangle” method (IJDefault only used if triangle produced positive values in any corner of the image). This mask was then cleaned to the single largest continuous piece (to remove speckles) then used to mask and automatically crop each channel. Conversion to data. Normalized, background subtracted, masked, cropped images were combined into a single image with each channel still separated. This image was converted into a data.frame. Within the data.frame, x and y coordinates were transformed into radial coordinates by finding the center of the masked area (approximation of the center of the colony with the radius converted from pixels into microns using the pixel ratio extracted from the original microscopy image. Binning. Radial bins were generated by dividing the image angularly by 256 radians and every 100 µm. Within each of these bins, the mean was taken for each channel, then the ratio between channels was calculated by subtracting one from the other and dividing them by the sum of both ((c1-c2)/(c1+c2)). Thus, values can range from 1/-1 indicating only the presence of one strain, to 0, indicating an exactly equal mix of the two strains. The proportion of overlapping bins was calculated by dividing the number of bins in the range -0.25:0.25 by the total number of bins. Area quantification. The individual channels of the normalized, background subtracted, masked, cropped images were thresholded using the imager::threshold() function to determine the area occupied by each species. Manual inspection of images indicated that for *P. aeruginosa* areas were underestimated and *K. pneumoniae* areas were over estimated. Therefore, all images were also manually quantified. For manual quantification, a combination of thresholding and area selection using the oval, wand or freehand selection was used. Comparison of manual and computational area quantification showed both methods produced similar results (Supplementary Figure 1).

**Statistics:** Analysis of variance was used to determine if there were significant differences in colony area between single and dual species biofilms. Data were grouped by timepoint and species, then Tukey's honest significant difference post-hoc test was used to determine p-values for specific comparisons. All tests were then pooled and a multiple-testing adjusted p-value was calculated using the R function p.adjust using the Benjamini-Hochberg correction. Separate analyses were calculated for the manual and computational area calculations, but the results from the manual quantification were considered more accurate so were used to draw conclusions about the data.

**A**

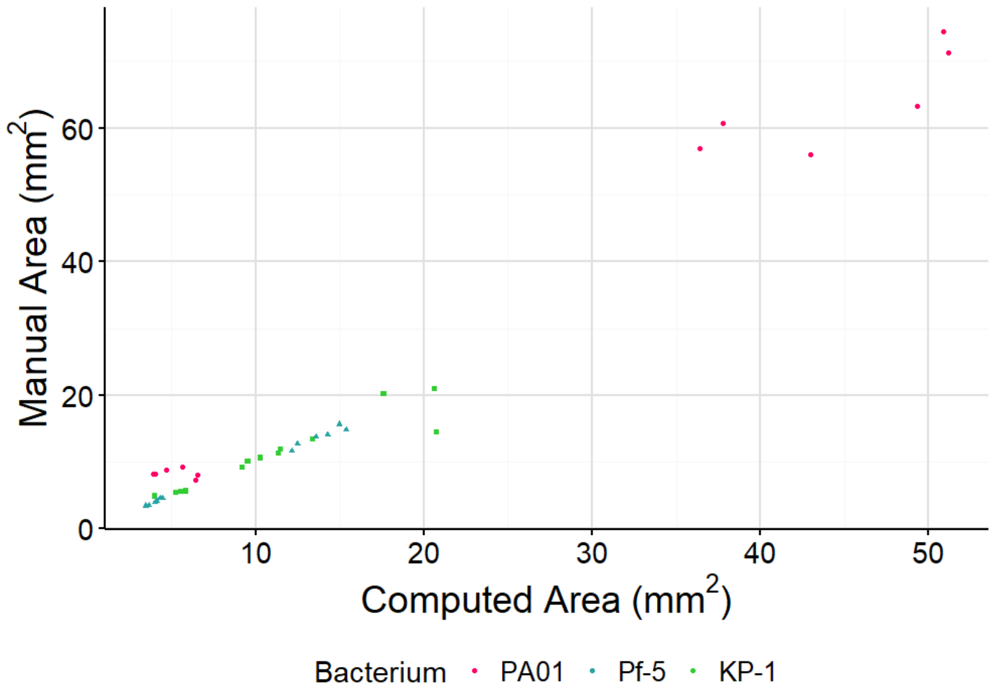

Manual Area (mm$^2$)
0
20
40
60
10
20
30
40
50
Computed Area (mm$^2$)
Bacterium
PA01
Pf-5
KP-1


**B**

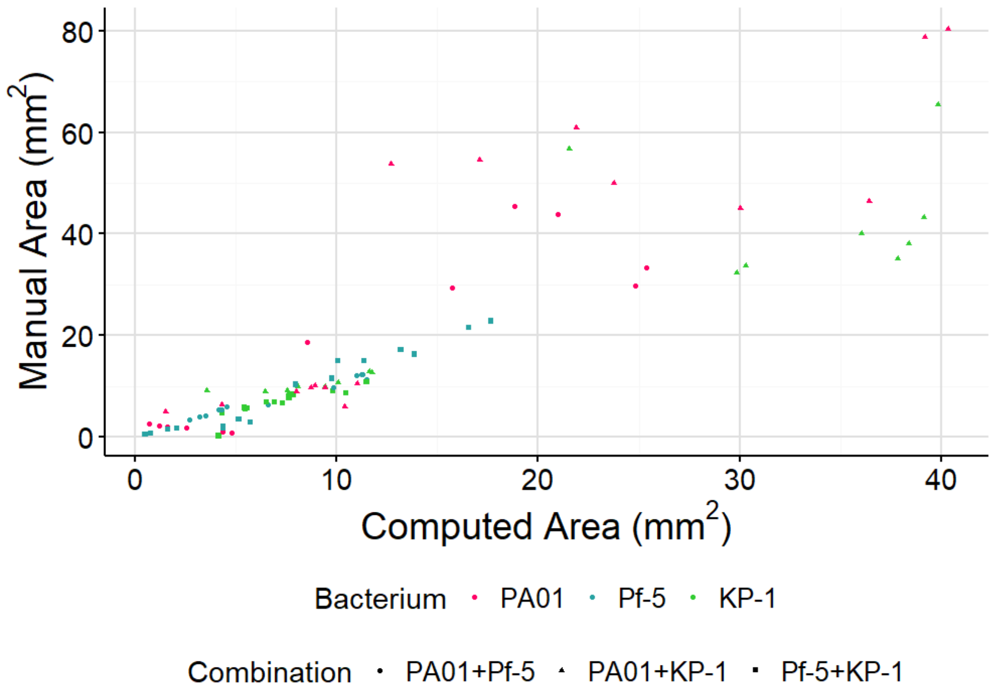

Manual Area (mm$^2$)
0
20
40
60
80
0
10
20
30
40
Computed Area (mm$^2$)
Bacterium
PA01
Pf-5
KP-1
Combination
PA01+Pf-5
PA01+KP-1
Pf-5+KP-1

**C**

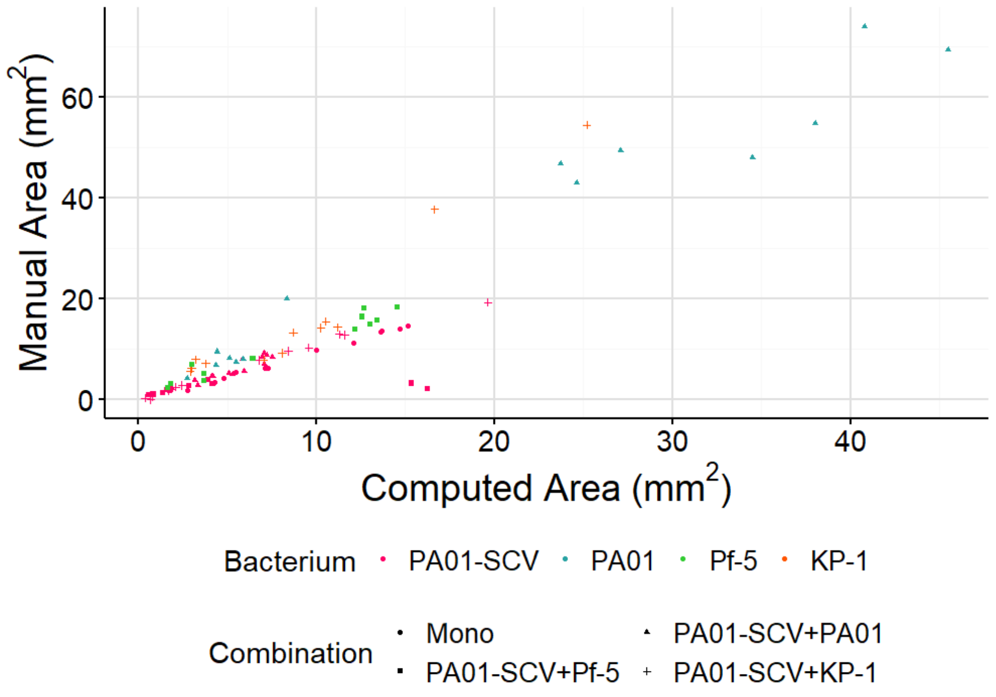
Manual Area (mm$^2$)
0
20
40
60
0
10
20
30
40
Computed Area (mm$^2$)
Bacterium
PA01-SCV
PA01
Pf-5
KP-1
Combination
Mono
PA01-SCV+PA01
PA01-SCV+Pf-5
PA01-SCV+KP-1

**D**

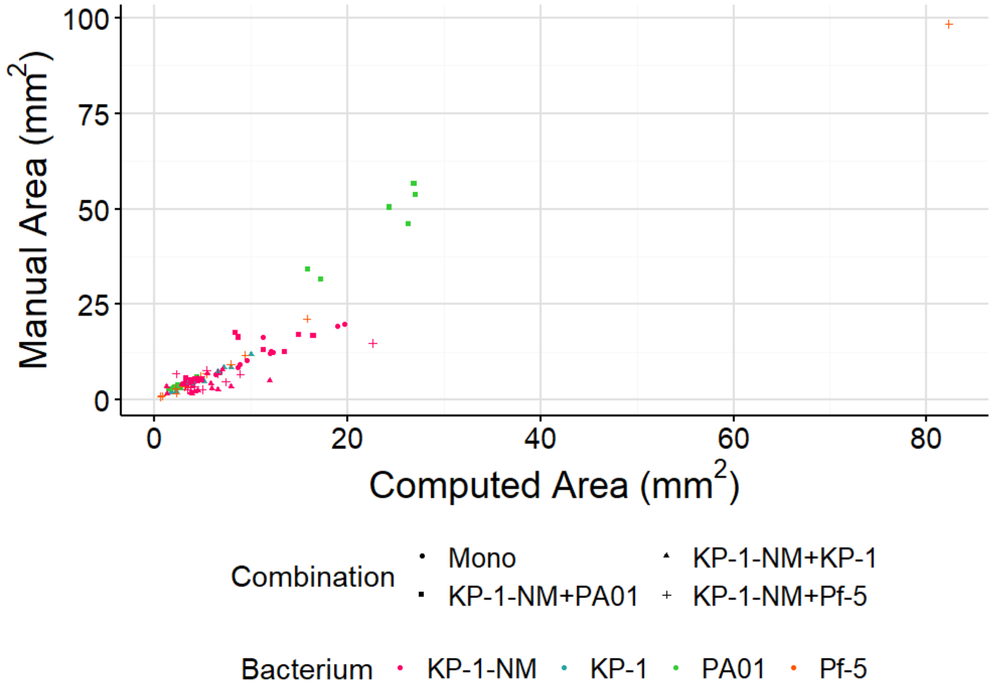
Manual Area (mm$^2$)
0
25
50
75
100
0
20
40
60
80
Computed Area (mm$^2$)
Combination
Mono
KP-1-NM+KP-1
KP-1-NM+PA01
KP-1-NM+Pf-5
Bacterium
KP-1-NM
KP-1
PA01
Pf-5

**E**

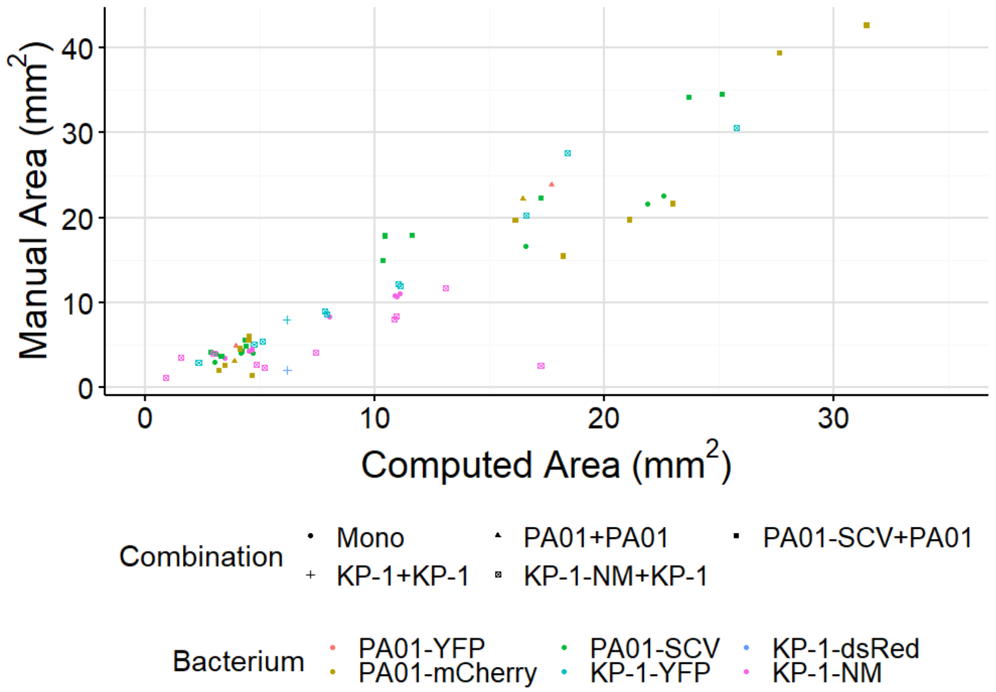


Supplementary Figure 1: Comparison of manual and computationally determined areas of colony biofilms of *P. aeruginosa*, *P. protegens* and *K. pneumoniae*. Single species (A), pair-wise co-cultures (B), co-cultures with the small colony variant (SCV) of *P. aeruginosa* (C), co-cultures with the non-mucoid variant (NMV) of *K. pneumoniae* (D) and colony biofilms grown on 0.6% agar (E) are compared separately. Each point represents a single replicate.

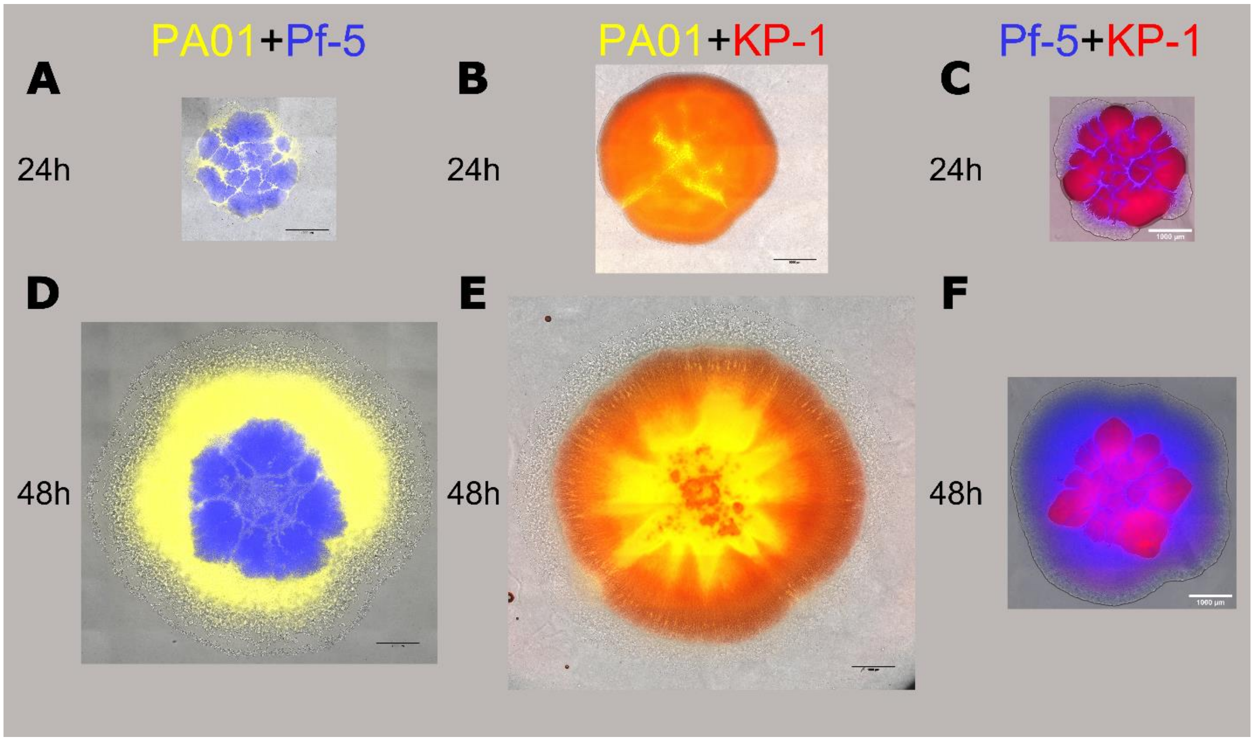


Supplementary Figure 2: Dual species colony biofilms consisting of a 1:1: mixture of fluorescently marked strains of *P. aeruginosa* PAO1 and *P. protegens* Pf-5 (A, D), *P. aeruginosa* PAO1 and *K. pneumoniae* KP-1 (B, E), and *P. protegens* Pf-5 and *K. pneumoniae* KP-1 (C, F), imaged after 24 and 48 h of growth. Strains are false coloured (PAO1, yellow; Pf-5, blue; KP-1, red). Fluorescent channels are overlaid with the brightfield channel. Image brightness was scaled manually for each channel in each image for visibility. The scale bar is 1000 µm. Note the visible colony in the brightfield of *P. aeruginosa* beyond what is visible in the fluorescent channels (D,E).

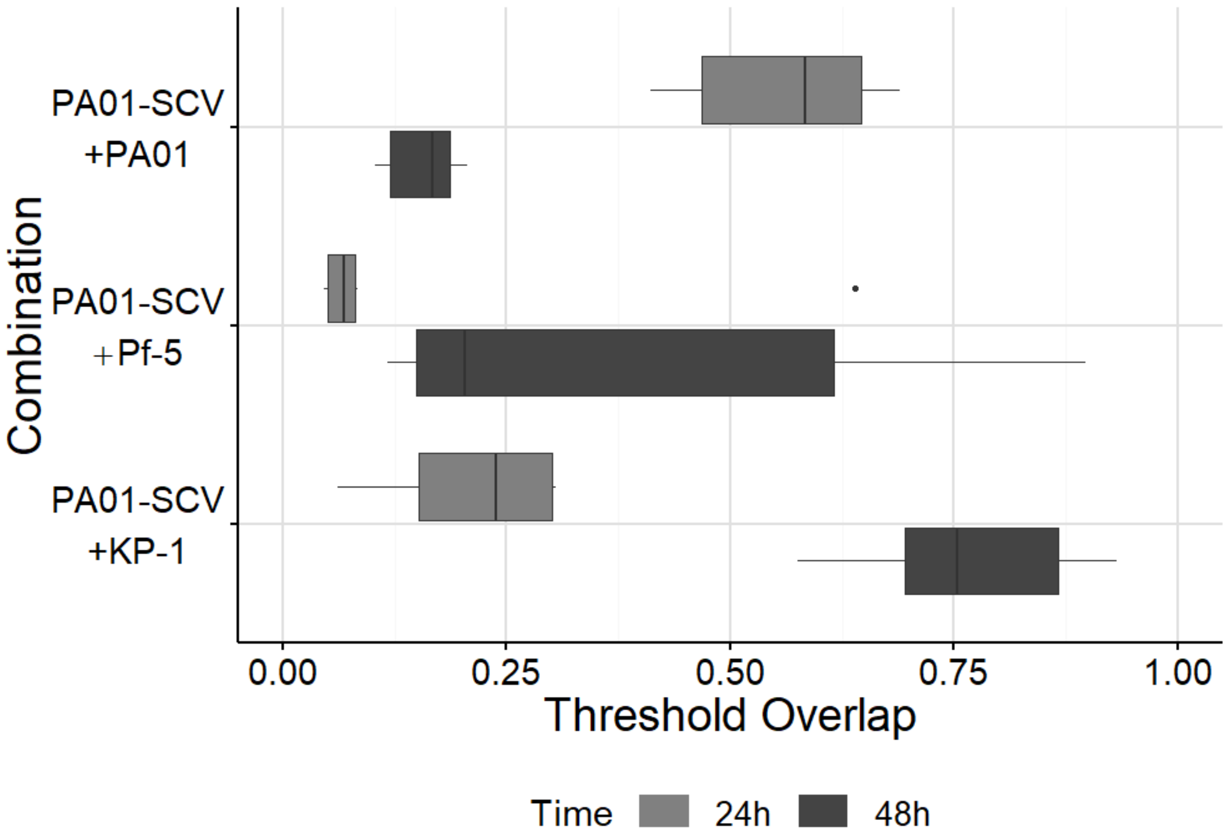


Supplementary Figure 3: Quantitative analysis showing co-localization of the different species, based on thresholded images of each channel rather than bin-based analysis. Overlap was calculated from raw images (n ≥ 3) which were split into separate channels then thresholded and the percentage of pixels with values of 1 in both channels used to define overlap. Compare to Figure 3 (G).

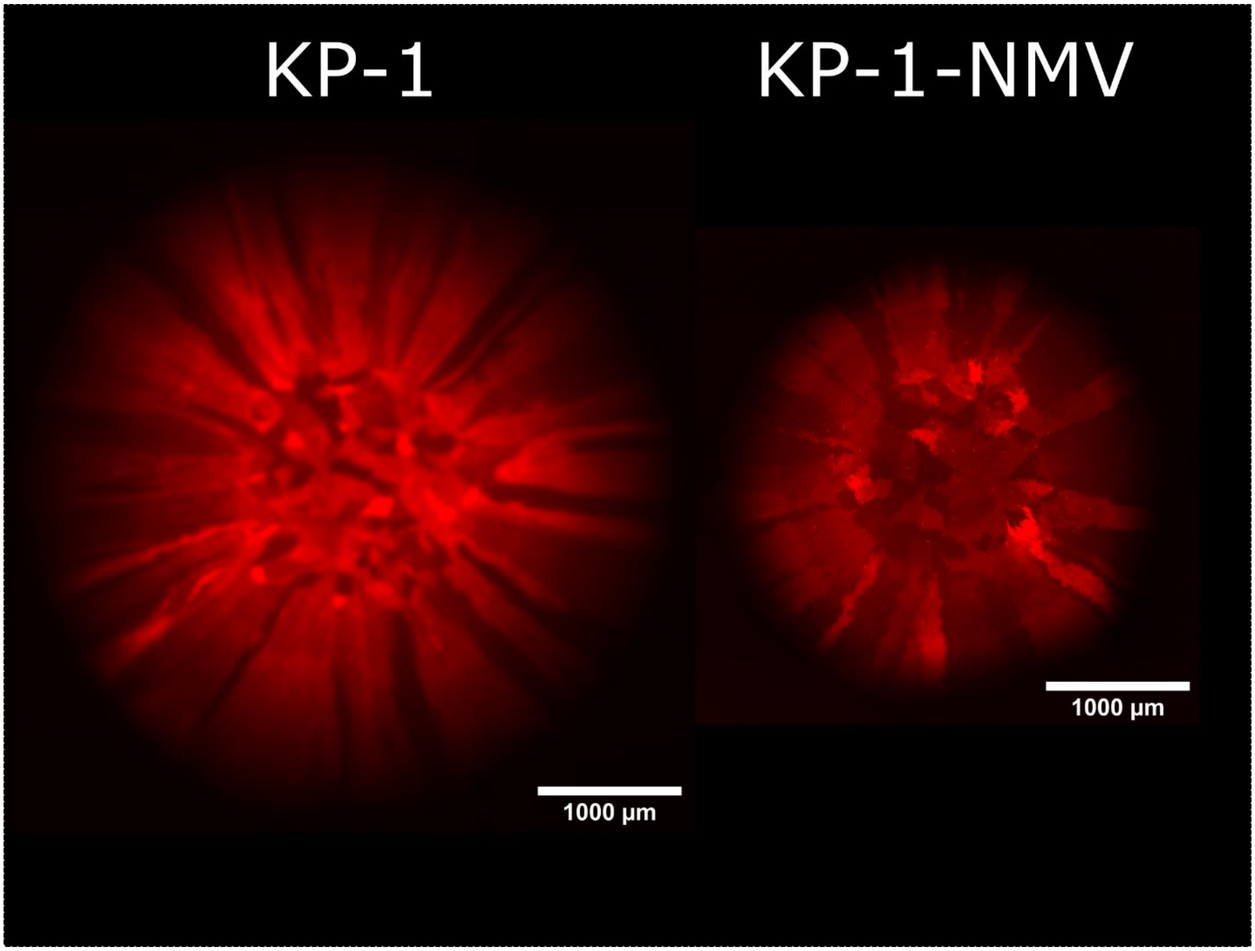


Supplementary Figure 4: Monospecies colony biofilms of *K. pneumoniae* KP-1 wild-type (left) and non-mucoid variant (NMV, right) expressing dsRed, imaged after 48 h of growth on 1.5% agar. Scale bar is 1000 µm. Note the different sectors due to differences in fluorescent protein expression. In the wild-type these boundaries are straight, but in the NMV note that there is a fractal quality. Sectors are visible due to variable expression of the dsRed fluorescent protein gene between founding cells.

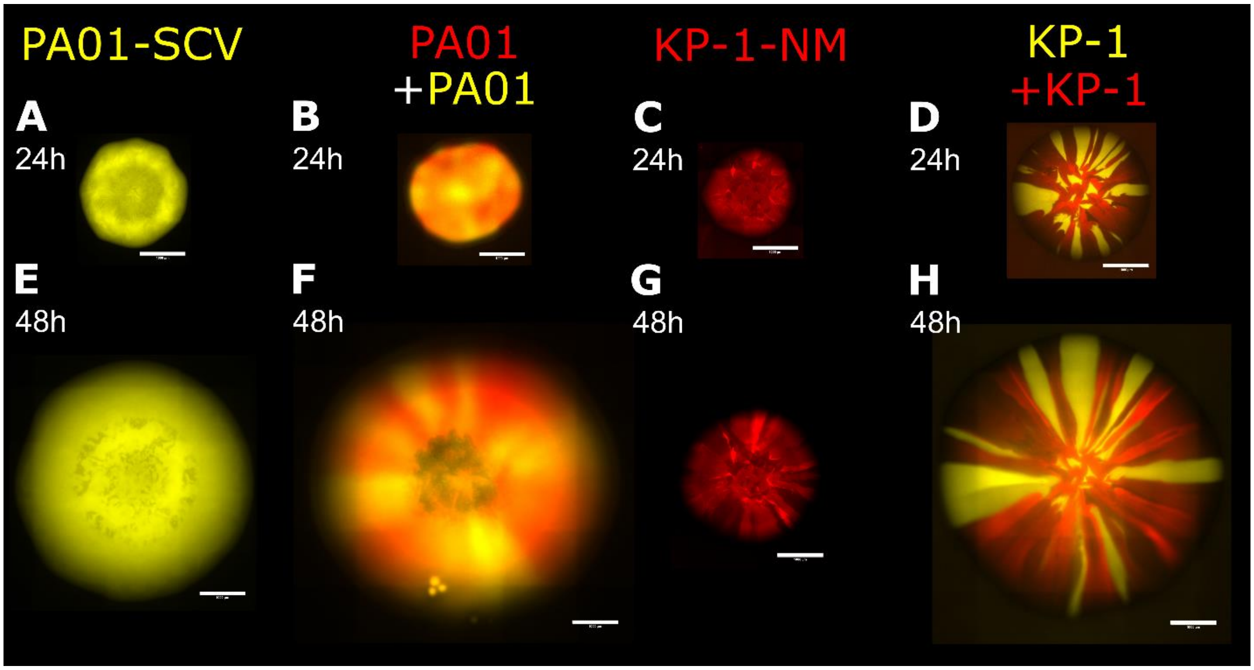


Supplementary Figure 5: Colony biofilms consisting of just *P. aeruginosa* small colony variant (SCV, A, E) or *K. pneumoniae* non-mucoid variant (NMV, C, G) or 1:1 mixtures of differently tagged wild-type strains (B, D, F, H), imaged after 24 and 48 h of growth on 0.6% agar. Scale bar indicates 1000 µm.

Supplementary Video 1: Video showing the leading edge of a colony biofilm comprised of *P. aeruginosa* PA01 strains expressing either YFP or mCherry fluorescent proteins. Frames are approximately 8 s apart, scale bar is 100 µm. Focus was maintained manually over the course of the 10 minutes. Each channel was separately subject to 'Bleach Correction' in ImageJ to correct for decreasing fluorescence over the course of the video capture.

Supplementary Video 2: Video showing the inner edge of a colony biofilm comprised of *P. aeruginosa* PA01 strains expressing either YFP or mCherry fluorescent proteins. Frames are approximately 8 s apart, scale bar is 100 µm. Focus was maintained manually over the course of the 15 minutes. Each channel was separately subject to 'Bleach Correction' in ImageJ to correct for decreasing fluorescence over the course of the video capture. The edge of the colony is located approximately at the top right corner. A line from the top right to bottom left of the field of view aims approximately towards the center of the colony.

Supplementary Table 1: Results from analysis of variance and Tukey's honest significance difference post-hoc test comparing colony biofilm areas. All tests were done on both computational and manually determined colony areas, then p-values were corrected for multiple hypothesis testing. Values below 0.05 are highlighted in blue.

| Time | Strain | Comparison | Computational | Manual |
|---|---|---|---|---|
| 24h | KP-1 | 0.6% KP-1-NM+KP-1-Mono | 0.332 | 0.388 |
| 48h | KP-1 | 0.6% KP-1-NM+KP-1-Mono | 0.349 | 0.090 |
| 24h | KP-1 | 1.5-0.6 | 0.092 | 0.084 |
| 48h | KP-1 | 1.5-0.6 | 1.81E-03 | 4.29E-05 |
| 24h | KP-1 | KP-1-NM+KP-1-Mono | 0.156 | 0.099 |
| 48h | KP-1 | KP-1-NM+KP-1-Mono | 0.081 | 0.571 |
| 24h | KP-1 | PA01-SCV+KP-1-Mono | 0.894 | 0.356 |
| 48h | KP-1 | PA01-SCV+KP-1-Mono | 0.995 | 0.299 |
| 24h | KP-1 | PA01+KP-1-Mono | 1.52E-01 | 5.35E-03 |
| 48h | KP-1 | PA01+KP-1-Mono | 8.04E-06 | 1.75E-05 |
| 24h | KP-1 | Pf-5+KP-1-Mono | 0.995 | 0.981 |
| 48h | KP-1 | Pf-5+KP-1-Mono | 0.349 | 0.696 |
| 24h | KP-1-NM | 0.6% KP-1-NM+KP-1-Mono | 0.332 | 0.166 |
| 48h | KP-1-NM | 0.6% KP-1-NM+KP-1-Mono | 0.341 | 0.228 |
| 24h | KP-1-NM | 1.5-0.6 | 0.821 | 0.141 |
| 48h | KP-1-NM | 1.5-0.6 | 0.555 | 0.228 |

| | | | | |
|---|---|---|---|---|
| 24h | KP-1-NM | KP-1-NM+KP-1-Mono | 0.314 | 9.54E-04 |
| 48h | KP-1-NM | KP-1-NM+KP-1-Mono | 0.042 | 4.23E-04 |
| 24h | KP-1-NM | KP-1-NM+PA01-Mono | 0.195 | 0.228 |
| 48h | KP-1-NM | KP-1-NM+PA01-Mono | 0.995 | 0.920 |
| 24h | KP-1-NM | KP-1-NM+Pf-5-Mono | 0.156 | 0.037 |
| 48h | KP-1-NM | KP-1-NM+Pf-5-Mono | 0.460 | 0.011 |
| 24h | KP-1/NM | 0.6% KP-1-KP-1-NM | 0.677 | 0.065 |
| 48h | KP-1/NM | 0.6% KP-1-KP-1-NM | 0.142 | 3.17E-04 |
| 24h | KP-1/NM | KP-1-KP-1-NM | 0.506 | 0.414 |
| 48h | KP-1/NM | KP-1-KP-1-NM | 0.974 | 0.414 |
| 24h | PA01 | 0.6% PA01-SCV+PA01-Mono | 0.055 | 5.93E-03 |
| 48h | PA01 | 0.6% PA01-SCV+PA01-Mono | 0.081 | 0.011 |
| 24h | PA01 | 1.5-0.6 | 0.304 | 0.522 |
| 48h | PA01 | 1.5-0.6 | 0.015 | 1.82E-03 |
| 24h | PA01 | KP-1-NM+PA01-Mono | 0.259 | 1.64E-03 |
| 48h | PA01 | KP-1-NM+PA01-Mono | 5.15E-03 | 0.066 |
| 24h | PA01 | PA01-SCV+PA01-Mono | 1.000 | 0.981 |
| 48h | PA01 | PA01-SCV+PA01-Mono | 0.216 | 0.639 |
| 24h | PA01 | PA01+KP-1-Mono | 0.269 | 0.996 |
| 48h | PA01 | PA01+KP-1-Mono | 0.023 | 0.761 |
| 24h | PA01 | PA01+Pf-5-Mono | 0.294 | 2.76E-05 |
| 48h | PA01 | PA01+Pf-5-Mono | 2.25E-03 | 2.53E-03 |
| 24h | PA01-SCV | 0.6% PA01-SCV+PA01-Mono | 0.092 | 0.334 |

| | | | | |
|---|---|---|---|---|
| 48h | PA01-SCV | 0.6% PA01-SCV+PA01-Mono | 0.216 | 0.653 |
| 24h | PA01-SCV | 1.5-0.6 | 0.314 | 0.537 |
| 48h | PA01-SCV | 1.5-0.6 | 7.85E-03 | 8.34E-03 |
| 24h | PA01-SCV | PA01-SCV+KP-1-Mono | 2.29E-03 | 4.30E-03 |
| 48h | PA01-SCV | PA01-SCV+KP-1-Mono | 0.964 | 0.981 |
| 24h | PA01-SCV | PA01-SCV+PA01-Mono | 0.358 | 0.939 |
| 48h | PA01-SCV | PA01-SCV+PA01-Mono | 0.152 | 0.015 |
| 24h | PA01-SCV | PA01-SCV+Pf-5-Mono | 4.37E-03 | 7.90E-03 |
| 48h | PA01-SCV | PA01-SCV+Pf-5-Mono | 0.177 | 1.87E-05 |
| 24h | PA01/SCV | 0.6% PA01-PA01-SCV | 0.677 | 0.023 |
| 48h | PA01/SCV | 0.6% PA01-PA01-SCV | 0.042 | 3.35E-04 |
| 24h | PA01/SCV | PA01-PA01-SCV | 0.995 | 2.71E-03 |
| 48h | PA01/SCV | PA01-PA01-SCV | 1.70E-05 | 1.15E-06 |
| 24h | Pf-5 | KP-1-NM+Pf-5-Mono | 0.195 | 0.029 |
| 48h | Pf-5 | KP-1-NM+Pf-5-Mono | 0.893 | 0.653 |
| 24h | Pf-5 | PA01-SCV+Pf-5-Mono | 0.773 | 0.639 |
| 48h | Pf-5 | PA01-SCV+Pf-5-Mono | 0.964 | 0.414 |
| 24h | Pf-5 | PA01+Pf-5-Mono | 0.995 | 0.622 |
| 48h | Pf-5 | PA01+Pf-5-Mono | 0.156 | 0.356 |
| 24h | Pf-5 | Pf-5+KP-1-Mono | 0.621 | 2.53E-03 |
| 48h | Pf-5 | Pf-5+KP-1-Mono | 0.786 | 0.497 |
